# Predicting intentional and unintentional task unrelated thought with EEG

**DOI:** 10.1101/764803

**Authors:** Adrien Martel, Mahnaz Arvaneh, Ian Robertson, Paul Dockree

## Abstract

Our attention seldom remains on a singular activity, instead veering off into thoughts unrelated to the task at hand. Studies adopting a component process view of off-task thought have begun to identify the underlying mechanisms and associated electrophysiological correlates underlying ongoing thought. In the present study, we developed subject-independent classification algorithms based on electroencephalographic (EEG) markers to discriminate on-task vs off-task as well as intentional vs unintentional off-task thought. To that end, spatio-temporal and spectral features extracted from EEG activity prior to reports of ongoing thought during a test of sustained attention were ranked according to their discriminative power. Using data collected from 26 participants, average classification accuracies of 83.4% and 71.6% were achieved using a regularized linear model for on-task vs off-task and intentional vs unintentional off-task thought, respectively. Our results identified gamma oscillations as the most discriminative feature to distinguish on-task from off-task states, and alpha synchronization as the most prominent feature when off-task states are engaged in deliberately rather than when experienced as arising spontaneously. Our work represents the first successful attempt at reliably discriminating the degree of intentionality experienced during task-unrelated thought and highlights the importance of recognizing the heterogeneous nature of off-task states.

## Introduction

Most often, the focus of our attention is not tied to the present moment or the immediate surroundings but rather engrossed in information generated from internal representations, such as autobiographical memories (Poerio et al., 2017; Smallwood et al., 2016; Smallwood & Schooler, 2006, 2015) or prospective thinking (Baird, Smallwood, & Schooler, 2011; Schooler et al., 2011; Spreng, Mar, & Kim, 2009; Stawarczyk, Cassol, & D’Argembeau, 2013). Ubiquitous both in daily life (Kane et al., 2007, 2017; Killingsworth & Gilbert, 2010; Poerio, Totterdell, & Miles, 2013; Risko, Anderson, Sarwal, Engelhardt, & Kingstone, 2012; Seli, Beaty, et al., 2018) and in the lab (McVay, Kane, & Kwapil, 2009; Smallwood, Davies, et al., 2004; Smallwood, Obonsawin, & Heim, 2003; Smallwood, O’Connor, Sudberry, Haskell, & Ballantyne, 2004; Varao-Sousa, Smilek, & Kingstone, 2018), these off-task experiences can vary along different dimensions such as content (Ruby, Smallwood, Engen, & Singer, 2013; Smallwood, Nind, & O’Connor, 2009; Smallwood & O’Connor, 2011), intrinsic or extrinsic constraints imposed on cognition (Christoff, Irving, Fox, Nathan Spreng, & Andrews-Hanna, 2016; Mills, Raffaelli, Irving, Stan, & Christoff, 2018), metacognitive awareness (Drescher, Van den Bussche, & Desender, 2018; Schooler, 2002; Schooler et al., 2011; Zedelius, Broadway, & Schooler, 2015) and degrees of intentionality (Martel, Arvaneh, Robertson, Smallwood, & Dockree, 2019; Robison & Unsworth, 2018; Seli, Carriere, & Smilek, 2014; Seli, Ralph, Konishi, Smilek, & Schacter, 2017; Seli, Ralph, Risko, et al., 2017; Seli, Risko, Smilek, & Schacter, 2016; Seli, Wammes, Risko, & Smilek, 2015). As mind-wandering research gains more traction in cognitive research and neuroscience, efforts have recently been put forth to resolve the two key issues stymying scientific progress, the ontological uncertainty due to the apparent heterogeneity of the phenomenon (Christoff et al., 2016, 2018; Seli, Kane, Metzinger, et al., 2018; Seli, Kane, Smallwood, et al., 2018; Wang et al., 2018) and the paucity of objective measures (Faber, Bixler, & D’Mello, 2018; Konishi & Smallwood, 2016; Martinon, Smallwood, McGann, Hamilton, & Riby, 2019).

Recently, views have emerged from both psychology (Seli, Kane, Smallwood, et al., 2018) and neuroscience (Wang et al., 2018) explaining off-task experiences as a family of states with overlapping experiential and neurocognitive features. The component process account (Smallwood & Schooler, 2015) explains the apparent heterogeneity of the off-task state as the result of varying contribution from discrete neurocognitive processes. Evidence for this view comes from studies showing that different facets of the off-task experience are a function of both shared and distinct neurocognitive profiles. For example, a recent study by the authors of the present paper (Martel et al., 2019) shows that one of the features of off-task thought, the degree of intentionality, can be understood as the interplay between bottom-up and top-down features of attention. More specifically, of the two processes that are assumed by contemporary accounts to be critical during off-task mental activity (Smallwood, 2013a), perceptual decoupling and executive control, the latter is thought to contribute exclusively to off-task thought when engaged in a deliberate manner. Given the limited capacity of attentional processing and that, by their nature, off-task thoughts are related to information absent from the immediate environment, it is assumed that these thoughts rely on the capacity by individuals to decouple attention from perceptual input to unfold (Smallwood, 2013b).

The electrophysiological dynamics of the perceptual decoupling process have been well studied and findings show that cortical processing of perceptual input is transiently dampened during off-task mental activity (Baird, Smallwood, Lutz, & Schooler, 2014; Barron, Riby, Greer, & Smallwood, 2011; Kam et al., 2011; Smallwood, Beach, Schooler, & Handy, 2008). This effect has proven to be robust across experimental paradigms and sensory modalities (Baldwin et al., 2017; Braboszcz & Delorme, 2011; Broadway, Franklin, & Schooler, 2015/4; Kam, Xu, & Handy, 2014; Martel, Dähne, & Blankertz, 2014; O’Connell et al., 2009). Generally speaking, perceptual decoupling during off-task states is associated in the EEG with a decrease of sensory and cognitive event-related potentials (ERPs (Hopfinger, Buonocore, & Mangun, 2000; Kam et al., 2011; Smallwood et al., 2008), an attenuation of inter-trial phase coherence (ITPC (Baird et al., 2014; Martel et al., 2019) and an upswing of low-frequency oscillations (Broadway et al., 2015/4; Jin, Borst, & van Vugt, 2019; Macdonald, Mathan, & Yeung, 2011; Martel et al., 2014; O’Connell et al., 2009). Taken together, these neurocognitive dynamics substantiate the notion that the perceptual decoupling process serves as an insulation mechanism, shielding cognition from competing input and allowing off-task thoughts to unfold (Smallwood, 2013a, 2013b).

In contrast, the process of executive control serves the important function of narrowing the focus of attention on information relevant to a specific train of thought, and in upholding off-task and task-relevant patterns of cognition with respect to the motivation and neuroanatomical structure of the individual (Golchert et al., 2017) or the demands of the task (Bernhardt et al., 2014; Smallwood, Brown, Baird, & Schooler, 2012). Although the role of perceptual decoupling is now well understood, the role of executive control in off-task states remains more elusive and a central question of the literature (Franklin, Mrazek, Broadway, & Schooler, 2013; McVay & Kane, 2010; Smallwood, 2013b; Smallwood & Schooler, 2006; Watkins, 2008), with evidence for its involvement in both off-task and task-related thought. The recent study led by the authors of the current work (Martel et al., 2019) investigating the intentionality dimension of off-task experiences found that spontaneous and deliberate forms of off-task states differ on several EEG features. Among these markers, alpha synchronisation, associated with perceptual inhibition (Foxe & Snyder, 2011) and internal attention (Benedek, Schickel, Jauk, Fink, & Neubauer, 2014; Cooper, Croft, Dominey, Burgess, & Gruzelier, 2003; Katahira et al., 2018), was found to be the most prominent marker of deliberate off-task states and interpreted as reflecting top down inhibition of irrelevant sensory input. Taken together, the neural data suggests that the level of intention during off-task states is determinant as to whether the experience is related to a failure of executive control or recruits similar mechanisms as task-related and off-task patterns of cognition.

Concurrent to these developments, the paucity of validated objective measures and the imperviousness of off-task thought to experimental manipulation has been a major methodological issue in mind-wandering research and still remains an open challenge (Smallwood & Schooler, 2015). Given the covert nature of off-task experiences, i.e. intrinsically driven and relating to information independent from the immediate environment, researchers have been bound to rely almost exclusively on experience sampling (ES) methods, that in turn rely on the introspective ability of individuals (Konishi & Smallwood, 2016).

Amongst ES methods, probe-caught (PC) and self-caught (SC) probes are the most widely used, with the difference that the former randomly interrupts the task while the latter requires a manual interruption before prompting participants to report on the content of their cognition prior to the interruption (Giambra, 1989; Weinstein, 2018). Both types have unique sets of drawbacks, for example SC probes typically only captures off-tasks states and may result in fewer reports if used exclusively (Schooler et al., 2011). Moreover, they represent a dual task, requiring participants to perform the experimental task and ongoing monitoring of their mental states which can influence the contents of cognition (Konishi & Smallwood, 2016). In contrast, PC probes allow for the collection of on-task periods, which is useful for juxtaposition with off-task states, but is prone to miss off-task experiences since it does not offer the possibility of manual reports and given that longer periods in-between probes has been shown to increase the likelihood of off-task reports (Seli, Carriere, Levene, & Smilek, 2013). In addition to these specific trade-offs, probes in general interrupt the flow of the task and offer no insight on whether off-task experiences continue after the probe, i.e. reactive mind-wandering (Allan Cheyne, Solman, Carriere, & Smilek, 2009), or whether the short break leads to focused attention (Ariga & Lleras, 2011; Helton & Russell, 2012). Moreover, the quality of subjective reports depends on numerous parameters such as an individual’s meta-cognitive ability (Schooler et al., 2011; Smallwood & Schooler, 2006), confidence (Seli, Jonker, Cheyne, Cortes, & Smilek, 2015) and motivation (Zedelius et al., 2015), and even the framing and format of the probe (Seli, Beaty, et al., 2018; Weinstein, De Lima, & van der Zee, 2018).

In light of these methodological shortcomings, researchers have begun to look for ways to overcome introspective bias or obviate the need for probes altogether, either by refining measures of ES, e.g. leveraging the heightened metacognitive ability of expert meditators (Ellamil et al., 2016; Fox et al., 2012; Girn et al., 2017), or via a *triangulation* process of subjective, behavioral and neural correlates of off-task thought (Konishi & Smallwood, 2016; Smallwood & Schooler, 2015). Previous work identified several measures modulated by off-task states, including behavioral measures such as speech patterns (Drummond & Litman, 2010), response time (Bastian & Sackur, 2013; Cheyne, Carriere, & Smilek, 2006; McVay & Kane, 2009; Stawarczyk, Majerus, Maquet, & D’Argembeau, 2011)), body movement (Carriere, Seli, & Smilek, 2013; Farley, Risko, & Kingstone, 2013; Seli, Carriere, Thomson, et al., 2014), reading speed (Franklin, Smallwood, & Schooler, 2011), and peripheral physiological measures such as eye movement (Frank, Nara, Zavagnin, Touron, & Kane, 2015; Reichle, Reineberg, & Schooler, 2010; Uzzaman & Joordens, 2011; Zhang, Miller, Sun, & Cortina, 2018), gaze (Faber et al., 2018; Huang, Li, Ngai, Leong, & Bulling, 2019; Mills, Bixler, Wang, & D’Mello, 2016), blinks (Grandchamp, Braboszcz, & Delorme, 2014; McIntire, McKinley, Goodyear, & McIntire, 2014; Smilek, Carriere, & Cheyne, 2010), pupil dilation (Franklin, Broadway, Mrazek, Smallwood, & Schooler, 2013; Konishi, Brown, Battaglini, & Smallwood, 2017; Smallwood et al., 2011), and neurophysiological measures (Baird et al., 2014; Broadway et al., 2015/4; Jin et al., 2019; Kam et al., 2011; A. Martel, Arvaneh, Taylor, Dockree, & Robertson, 2017; Martel et al., 2019; O’Connell et al., 2009; Smallwood et al., 2008).

The next step in the triangulation approach after the collection of various measures is to integrate these into machine learning models to obtain a continuous index of attention. Fortuitously, brain-computer interfaces (BCIs) have garnered much attention in the recent past due to the attractive possibility of identifying and monitoring covert mental states based on brain activity in real-time (Blankertz et al., 2016, 2010; Venthur, Blankertz, Gugler, & Curio, 2010). Besides the communication and control application areas, BCIs are well suited as a potential research tool (Blankertz et al., 2016; Brunner et al., 2015; Vansteensel, Kristo, Aarnoutse, & Ramsey, 2017), for example by decoding brain signals and accurately discriminating between periods of off-task thought and on-task activity. Importantly, a BCI able to continuously index attention would help resolve the, yet unknown, temporal dynamics of off-task experiences; A knowledge gap that forces researchers to rely on estimates. For example, the 10 sec period prior to a probe associated with the reported attentional state used in the present work is an estimate based on prior studies and findings on the correlates of perceptual decoupling (Baird et al., 2014; Baldwin et al., 2017; Christoff, Gordon, Smallwood, Smith, & Schooler, 2009; Henríquez, Chica, Billeke, & Bartolomeo, 2016; Martel et al., 2014; O’Connell et al., 2009). Moreover, although the brain regions and neurophysiological correlates of processes in charge of maintaining off-task states are relatively well known (Andrews-Hanna, Smallwood, & Spreng, 2014; Bernhardt et al., 2014; Christoff et al., 2009) and given the assumption that off-task thought occurs in stages (Girn et al., 2017; Smallwood, 2013a) a continuous index of attention could provide valuable information on the neurocognitive processes and neuroanatomical structures involved in the onset of off-task experiences.

Research in that domain is inching ever closer to a general index of attention with studies identifying predictors of off-task states from oculometric data during breath-counting (Grandchamp et al., 2014) and reading (Faber et al., 2018), as well as from EEG data during a SART (Jin et al., 2019; Kawashima & Kumano, 2017; A. Martel et al., 2017) and simulated driving (Baldwin et al., 2017; Guo, Pan, Zhao, Cao, & Zhang, 2018). In of these studies, (Jin et al., 2019) used support vector machines to build a task-independent classifier able to discriminate on-task from off-task states across a SART and a visual search task. Interestingly, the authors identified alpha synchronization, which we previously found to be characteristic of intentional off-task states (Martel et al., 2019), as the most predictive feature, suggesting that at least some of the off-task states collected were of the deliberate kind. Although demonstrating the feasibility of a general classifier of on- vs off-task states is a vital achievement towards an accurate index of attention, off-task states are not homogenous with contrasting experiences relating to varying neurocognitive profiles, likely to produce distinct neural signatures.

Therefore, in the present work we make the first attempt at reliably classifying different degrees of intentionality when individuals experience off-task mental activity. To that end, we first compared spatio-temporal and spectral features extracted a priori based on previous findings and classified the neural data of on-task and off-task periods collected with PC probes during a SART to confirm previous findings, before juxtaposing periods of deliberate (dTUT) and spontaneous task-unrelated thoughts (sTUT) collected with SC probes.

## Methods

The neural data analysed here was collected in the context of a prior study. For the sake of brevity this section will give an overview of the experiment conducted and elaborate on the preprocessing and off-line classification. For a detailed explanation of the experimental setup please refer to (Martel et al., 2019).

### Setup

Twenty-six participants (12 females, age M: 25, SD: 4.3), with no known neurological disorder or psychiatric illness, performed a modified SART during which digits from 1 to 9 were sequentially presented centrally for 250 ms with an ISI of 2315 ms. Participants were instructed to respond to each digit except for the target ‘3’ and to interrupt the task when they became aware of an off-task thought, i.e. trigger an SC probe. PC probes pseudo-randomly interrupted the task after 28, 42, 56, 76, 83, 97 or 111 sec relative to the appearance of the last probe. Probes prompted participants to categorize their ongoing thought prior to the interruption as being (1) on-task, (2) concerning the task, (3) distracted by sensations, (4) deliberate task-unrelated thoughts (dTUT) or (5) spontaneous task-unrelated thoughts (sTUT). These five categories are an attempt at discretizing the spectrum of conscious states with regard to off-task thought (Seli et al., 2016; Smallwood & Schooler, 2015; Stawarczyk et al., 2011) allowing for a more granular resolution of off-task experience and avoiding the dichotomous assumption of most mind-wandering studies that deem all inattentive states to be mind-wandering. Participants were given extensive instructions on how to categorize thoughts and given a quiz with prototypical thoughts that they had to associate with each category before the start of the experiment. All procedures were reviewed and approved by the Trinity College Dublin ethics committee.

### Data

64-channel EEG data was acquired at 512 Hz using a BioSemi ActiveTwo system (biosemi.com) placed according to the international 10-20 system. All data were processed, analyzed and visualized through Matlab R2017a (Mathworks) with the help of custom written scripts and the following toolboxes: EEGLAB^93^, Fieldtrip^94^ and the BBCI toolbox (http://www.github.com/bbci/bbci_public). The pre-processing of the EEG data was kept rudimentary to simulate the input the BCI would receive and be able to process in real-time. Hence, the EEG data was subjected to a 0.5 Hz zero-phase high-pass filter, to remove slow trends, downsampled to 250Hz and robustly re-referenced with the PREP pipeline (Bigdely-Shamlo, Mullen, Kothe, Su, & Robbins, 2015). The EEG was segmented into epochs according to SART trials, i.e. appearance of the digit, with epochs spanning −500 ms to 2400 ms relative to stimulus onset. The reported attentional state was estimated to span four SART trials prior to the appearance of the probe (see (Martel et al., 2019)). The on-task condition (PC_OT) consisted of the on-task reports from PC probes and the off-task condition (PC_MW) of both deliberate and spontaneous reports from PC probes. The SC probes provided the intentional (SC_dTUT) and unintentional (SC_sTUT) off-task conditions from the corresponding reports. This configuration resulted in a total of 768 PC_OT, 704 PC_MW, 2434 SC_dTUT and 2358 SC_sTUT epochs across all the participants. To avoid biasing the classifier, compared conditions were balanced by randomly subsampling from the condition with more epochs for a total of 704*2 epochs for the PC_OT/MW comparison and 2358*2 epochs for the SC_dTUT/sTUT comparison.

### Feature extraction

For the identification of the mental states of interest, informative features must be extracted from the recorded EEG signals to form discriminative sets of features. Brain signals recorded by EEG are either of the exogenous (also called evoked) type, e.g. ERP components elicited in response to task stimuli, or of the endogenous (also called spontaneous) type such as the tonic alpha synchronization seemingly characteristic of deliberate off-task thoughts. Consequently, two types of features were leveraged in the present work: spatio-temporal features and spectral features. Spatio-temporal features relate to the spatial, i.e. the location of the channels on the scalp, and temporal, i.e. the fluctuations of potential over time for each channel, domain of the EEG signal. Conversely, the spectral features relate to the frequency domain, e.g. increased alpha synchronization during dTUT. For the classification of the conditions, spatio-temporal and spectral features were concatenated to build the feature vectors. Combining spatio-temporal and spectral features is commonly used in the design of BCIs to increase the amount of evidence for the training of the classifier (Dornhege, Blankertz, Curio, & Müller, 2003).

We performed iterative runs of feature extraction and classification, gradually adding and testing features and selecting for the most discriminative ones with the aim of surpassing a classification accuracy threshold of 70%, above which the classifier was considered as rigorous (A. Kübler, Mushahwar, Hochberg, & Donoghue, 2006; Andrea Kübler, Neumann, Wilhelm, Hinterberger, & Birbaumer, 2004). This procedure allowed us to test a prioris, constrain the search space and draw conclusions about the neurophysiological correlates most characteristic of each attentional state. We began with testing a priori chosen spatio-temporal features based on our previous findings (Martel et al., 2019) against those identified with an automated heuristic algorithm to identify the 5 most discriminative spatio-temporal features over all channels for each condition pair (See next section for more details). Subsequently, we combined these 5 features with a priori chosen spectral features over 3 sets of channels (averaged over fronto-central, parieto-occipital and all channels) for each condition-pair, e.g. delta+theta for PC_OT/MW, and tested 2 time-segments (whole epoch and second half) against each other (See Figure 2). Finally, for each condition-pair we selected the time-segment and 5 spatio-temporal features that returned the best classification accuracies and tested for the most discriminative spectral features by extracting power from the delta, theta and alpha range, as well as 16 overlapping bands from 15 to 100 Hz (See next section for more details).

### Spatio-temporal & spectral features

For extracting spatio-temporal features, a separability index was calculated for each condition-pair. The discriminability across conditions (or classes), e.g. between PC_OT and PC_MW, was determined with the signed point-biserial correlation coefficient (signed r^2^; (Blankertz, Lemm, Treder, Haufe, & Müller, 2011)). Subsequently, the EEG data of different time-intervals were ranked according to the highest measure of separability, i.e. signed r^2^ values, and added to the feature vector. For the a priori selected spatio-temporal features the search heuristic was constrained temporally and spatially according to previously known differences between conditions, e.g. P1 (80-110 ms relative to stimulus onset), P2 (200-300 ms) and P3 (270-380 ms) amplitude peak time segment for the condition pair SC_dTUT/sTUT (Martel et al., 2019), The number of spatio-temporal features was limited to the 5 per condition-pair with the highest r-values.

Similar to the feature extraction performed for the spatio-temporal features, the spectral features were first extracted according to prior knowledge. The a priori extracted spectral features were the power of delta + theta (1-7 Hz) band for PC_OT/MW and alpha (8 - 14 Hz) band for SC_dTUT/sTUT, since these bands exhibited statistical differences for their condition-pairs (Martel et al., 2019). We also found that the tonic activity for the delta+theta and alpha band occurred after the offset of the evoked activity (approximately 1200 msec relative to stimulus onset) and appeared to be localised in some cases, e.g. fronto-centrally for the theta power. To determine whether the choice of the time-interval for the band power calculation influenced the classification accuracy, the band power was calculated both for the entire trial (0 to 2400 ms relative to stimulus onset) and for the second half of the trial (1200 to 2400 msec). Similarly, to determine whether the spatial distribution of neural oscillations had an impact on classification results, the band power was calculated and averaged over all channels and over 2 subsets; fronto-central with 18 channels and parieto-occipital with 19 channels (see Figure 1). Hence, spectral features were extracted from the 3 sets of channels resulting in feature vectors of 5 (spatio-temporal features) + 1 (spectral features) = 6 features for each channel set.

**Figure 1.**
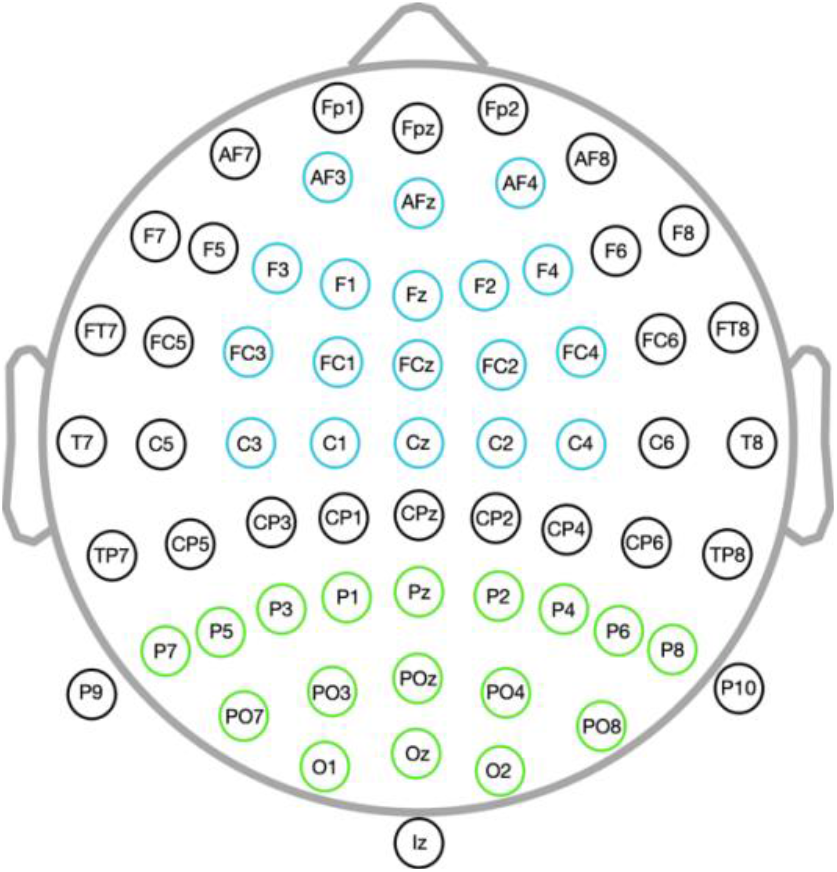
Channel location and nomenclature from the 10-20 international system of the 64-channel scalp Biosemi cap with the channels in blue belonging to the fronto-central selection and in green the channels belonging to the parieto-occipital selection.

Finally, for the last run, we divided the frequency domain into a total of 19 bands to sweep the frequency domain in which neural oscillations operate to identify additional discriminative spectral features. The first 3 bands spanned 1-3 Hz for the delta range, 4-7 Hz for the theta range and 8-14 Hz for the alpha range. The remaining 16 bands spanned 10 Hz and divided the frequency domain equally from 15 to 100 Hz with a 5 Hz overlap. The averaged power of each of the 19 bands were added individually as features resulting in feature vectors composed of 5 (spatio-temporal) + 19 (spectral features) = 24 features for each of the 3 sets of electrodes.

To estimate the band power for spectral features, the bandpower function of Matlab was used for convenience as it returned similar results to the Welch method (splitting the intervals into 8 segments with 50% overlap) or averaging the power extracted via wavelet decomposition. The bandpower function estimates the power by applying a Hamming window and using a periodogram of the same length as the input vector. Baseline normalization was achieved by dividing the power calculated for each epoch by the averaged power of the epoch formed from the trial preceding the succession of the four SART trials associated with a condition, i.e. the fifth SART trial when counting back from the probe.

### Feature Selection and Classification

For off-line classification, Linear Discriminant Analysis (LDA) with shrinkage of the covariance matrix was applied (Blankertz et al., 2011). LDA is a common classification method used in BCI experiments and has been shown to be effective in terms of minimizing misclassification errors (Duda, Hart, & Stork, 2012). Although more sophisticated classification methods (e.g. neural networks) have been shown to provide slightly better classification accuracies, the advantages over linear methods are outweighed by the higher computational costs and the tendency to overfit (Garrett, Peterson, Anderson, & Thaut, 2003).

Classification accuracies were computed under a stratified 10×10-fold cross-validation procedure. To remove redundant features and determine the best features in terms of classification accuracy, the features were ranked according to their classification accuracy by applying the ten-fold cross-validation with LDA to each feature individually over 100 iterations. The features with highest average classification accuracy were subsequently combined incrementally according to their rank. A series of classifications were then performed starting with the best feature and incrementally adding each feature according to their rank (first classification with the best feature, second with the two best features and so on).

To get better insight into the performance of features and to determine whether features differed in their discriminative power according to spatial locations three different channel arrangements were considered: (1) fronto-central (F1-3, Fz, FC1-4, FCz, C1-4, Cz, AF3-4, AFz), (2) parieto-occipital (P2-8, Pz, PO3-4, PO7-8, POz, O1-2, Oz) (3) all channels (see Figure 1).

Lastly, to provide an overview of the findings and to help with the interpretation of the most discriminative features, the average band power was extracted for the alpha (8 - 14 Hz), beta (15 - 40 Hz), lower gamma (40 - 70 Hz) and high gamma (80 - 100 Hz) bands over all non-boundary channels and for both condition-pairs. The data was subsequently binned according to percentiles (5 bins each representing 20% of the data) and plotted into a bivariate histogram depicting the proportional distribution of conditions against percentile bins.

## Results

### Probe-caught on-task vs off-task (PC_OT/MW) classification results

Given that the automated heuristic algorithm of the bbci toolbox function consistently returned better classification results, a priori selected time-intervals were dismissed in favor of the heuristic for the extraction of the 5 spatio-temporal features. The 5 spatio-temporal feature sets with the highest discriminability identified by the automated heuristic algorithm from the discriminability matrix of signed r^2^-values over all channels are shown in Figure 2. Two of the five identified time-intervals are of particular interest because the high discriminability they exhibit corroborate findings from our previous analysis of data (Martel et al., 2019). The time interval 120 to 144 msec and 336 to 396 msec corresponds to the P1 ERP and the P3 time-interval respectively, which we found to differ significantly in the ERP analysis. Although these two time intervals resemble the segments we chose initially, 80-110 ms for P1 and 270-380 for P3, it may be the case that the use of filters during our previous analysis, e.g. ERP averaging, introduced a temporal shift, which in turn might explain why features chosen by the heuristic differ and performed consistently better in the classification.

**Figure 2:**
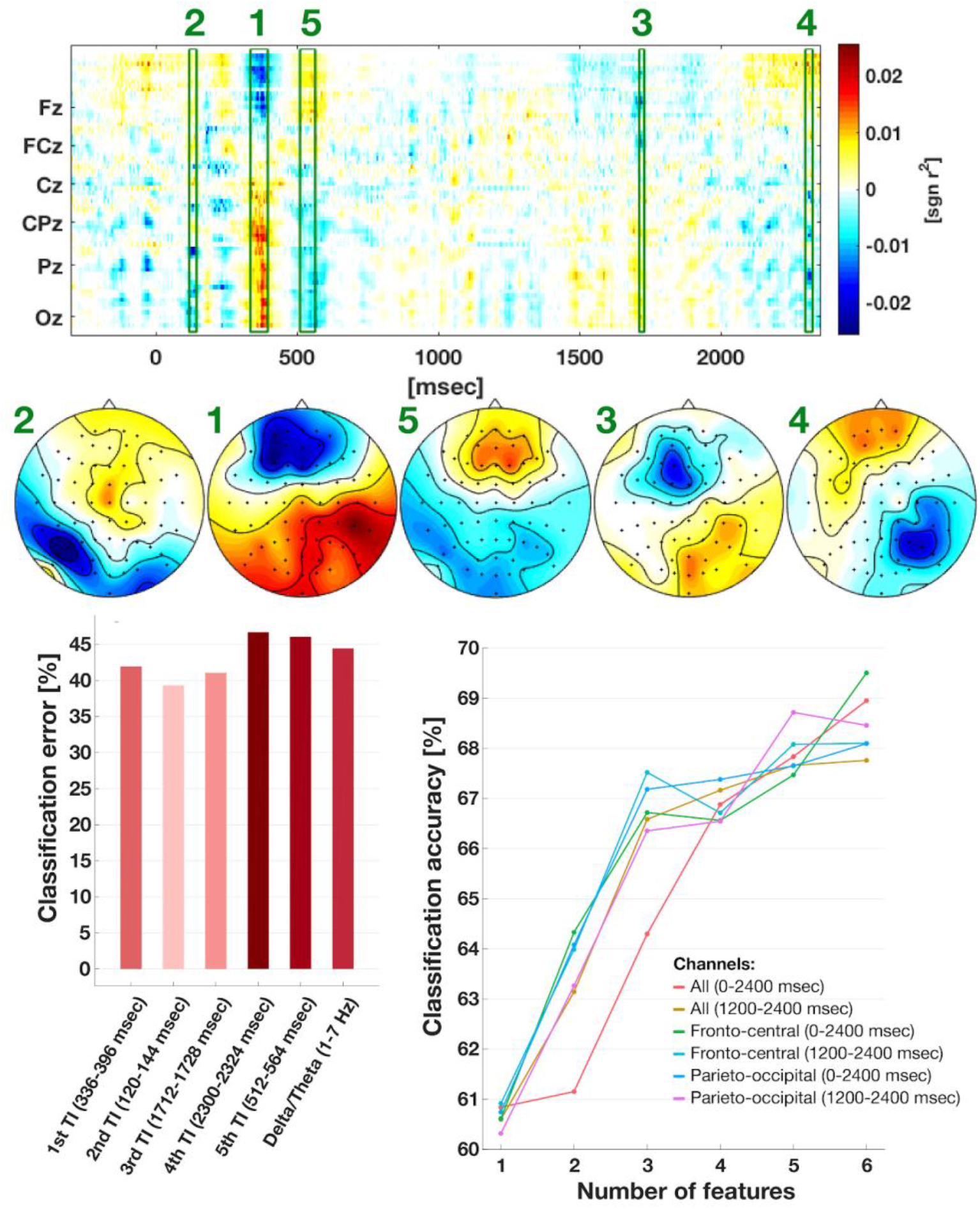
Spatio-temporal and Spectral classification of PC_OT/MW. TOP: Visualization of the signed r^2^ matrix. For the spatio-temporal features, signed r^2^-values of PC_OT minus PC_MW were calculated and depicted as a colour-coded matrix. MIDDLE: The scalp topographies of the averaged r^2^-values are depicted for the corresponding time intervals. Interestingly, the heuristic algorithm of the bbci toolbox identified the P1 (120 - 144 msec) and P3 (336 - 396 msec) time-intervals as being discriminative. The a priori chosen delta+theta band (1 - 7 Hz) was also included as a feature. BOTTOM LEFT: Averaged contribution to the classification error of each of the 5 spatio-temporal features and the 1 spectral feature (1 - 7 Hz). Intensity of the colour denotes the contribution to the classification error with lighter colours corresponding to more discriminative features. BOTTOM RIGHT: Classification accuracies (averaged from 100 cross-validations) of iteratively added features ranked according to their discriminative power. The spectral features included in the classification are shown for the band power calculated over two intervals (0 - 2400 and 1200 - 2400 msec) for three sets of channels (fronto-central, parieto-occipital and all channels). The x-axis depicts the number of features considered in the classification. The y-axis depicts the classification accuracy in percentage. The highest classification accuracy (69.4 %) was achieved with 6 feature sets from the fronto-central channels (i.e 5 temporal features and one band power delta+theta feature).

The first classification with 5 spatio-temporal features and 1 spectral feature returned a maximum average classification accuracy of 69.4% for the fronto-central channels and the band power calculated for delta+theta band over the entirety of the trial (0 - 2400 msec). The delta+theta band power measured over the entirety of the trial as opposed to the second half improved the classification of an average of 1.2% (Figure 2, bottom right panel). Consequently, the spectral features were computed on the entire trial for the remainder of the analyses. It is worth noting that the general trend for the classification accuracy shows an increase as a function of the number of incrementally added features, suggesting that it is likely to further improve with the addition of more features.

Building on these results the second series of classifications included the 15 spectral features extracted from the entire duration of the trial. The results of the classification can be taken from Figure 4.3. The overall highest classification accuracy of 83.4 % was obtained with 13 features, each averaged across all channels. The classification for the fronto-central channels did not improve substantially past the addition of a fourth feature (75.3 %) and peaked at 76.4 % for 16 features. The parieto-occipital channels fared the worst, peaking at 7 features with a classification accuracy of 73.5%. The most discriminative bands were found in the beta band (25 - 35 Hz), lower gamma band (50 - 70 Hz) and the higher gamma band (80 - 100 Hz). Crucially, the classification could be considered rigorous (above 70 %) for all three channel sets with a small number of features for parieto-occipital channels and with only one feature (spectral feature 55 - 60 Hz) for the other two sets of channels.

### Self-caught deliberate vs spontaneous TUT (SC_dTUT/sTUT) classification results

The classification applied on the SC-dTUT/sTUT followed the same procedure as described for PC-OT/MW with 3 spatio-temporal features, with a heuristic search over the signed r^2^-values matrix, and the same 16 spectral features. The results of the iterative classification is depicted in Figure 4. The highest classification accuracy of 71.6 % was obtained for 17 features for fronto-central channels with spectral features extracted from the entirety of the trial (0 - 2400 msec). The classification with spectral features extracted over the entire trial and all channels with 12 features reached 69.9%, almost crossing the BCI usability threshold. Interestingly, amongst the most discriminative features were the spatio-temporal features corresponding to the P2 ERP and the alpha band, corroborating previous findings (Jin et al., 2019; Martel et al., 2019).

### Percentile Distribution of Band Power

To facilitate the interpretation of the discriminative features identified, the probability density distribution of the number of epochs with respect to the alpha, beta, low and high gamma power is depicted in Figure 5. The most discriminative spectral features for PC_OT/MW condition pair were beta, low and high gamma power. The percentile distribution indicates that the PC_OT condition is characterised by an increased number of high-power low and high gamma activity relative to PC_MW (see Figure 5, left). Additionally, epochs with mid-power beta activity were underrepresented for the PC_MW condition. For SC-dTUT/sTUT, a higher distribution of epochs with high alpha and beta activity can be observed for dTUT (see Figure 5, right).

## DISCUSSION

Our study provides evidence that off-task states with different experiential features can be reliably differentiated on the basis of spatio-temporal and spectral features of the EEG. Replicating prior findings (Jin et al., 2019), we found that on-task (PC_OT) and off-task (PC_MW) states can be discriminated with an accuracy of 83.4% using linear models. More importantly, for the first time we show that intentional (dTUT) and unintentional (sTUT) off-task states can be differentiated with an accuracy of 71.6%. Crucially, P2 amplitude and alpha synchronization were amongst the most discriminative features, corroborating prior studies (Jin et al., 2019; Martel et al., 2019) and further establishing these features as markers of intentional off-task experiences.

Our classification results show that gamma band features were consistently amongst the most discriminative features, in particular for the on-task vs off-task (PC_OT/MW) condition pair. Interestingly, the bivariate histogram of the distribution of the number of epochs according to their power suggest that PC_OT is characterised by a greater number of epochs with high gamma activity (both low and high gamma). The same effect was found, albeit to a lesser extent, for SC_dTUT/sTUT. Gamma oscillatory activity (> 40 Hz) is known to be involved in a wide range of fundamental cognitive functions such a spatial attention (Kaiser & Lutzenberger, 2005), and memory (Jensen, Kaiser, & Lachaux, 2007). Gamma is generally thought to underlie feature binding (Treisman & Gelade, 1980) and reflect cortical activation (Merker, 2013). Within the context of attention, faster gamma oscillations are linked with processing of attentional processing (Brovelli, Lachaux, Kahane, & Boussaoud, 2005; Fries, Reynolds, Rorie, & Desimone, 2001; Womelsdorf & Fries, 2006) and maintenance of internal representations (Herrmann, Munk, & Engel, 2004). Moreover, the maintenance of task-related patterns required for sustained attention is believed to be facilitated by localised gamma activity in task-relevant brain areas (for a review see Clayton, Yeung, & Cohen Kadosh, 2015). Hence, an interesting aspect of the present findings is the contribution of the gamma activity to the successful classification of PC_OT/MW (Figure 3) and of SC_dMW/sMW with fronto-central channels (Figure 4), and the presence of more high-powered gamma epochs for the on-task condition (Figure 5). Considering that on-task states relative to off-task states, and dTUT relative to sTUT presumably rely on similar top-down mechanisms to maintain task-related and off-task patterns of cognition, it is supposable that gamma activity reflects executive control processes. Echoing this possibility, prior studies have associated gamma power changes with focused attention in experienced meditators (Braboszcz, Cahn, Levy, Fernandez, & Delorme, 2017; Cahn, Delorme, & Polich, 2010, 2013; Hinterberger, Schmidt, Kamei, & Walach, 2014; Lutz, Greischar, Rawlings, Ricard, & Davidson, 2004), and with internally directed attention, e.g. mental arithmetic (Ishii et al., 2014).

**Figure 3.**
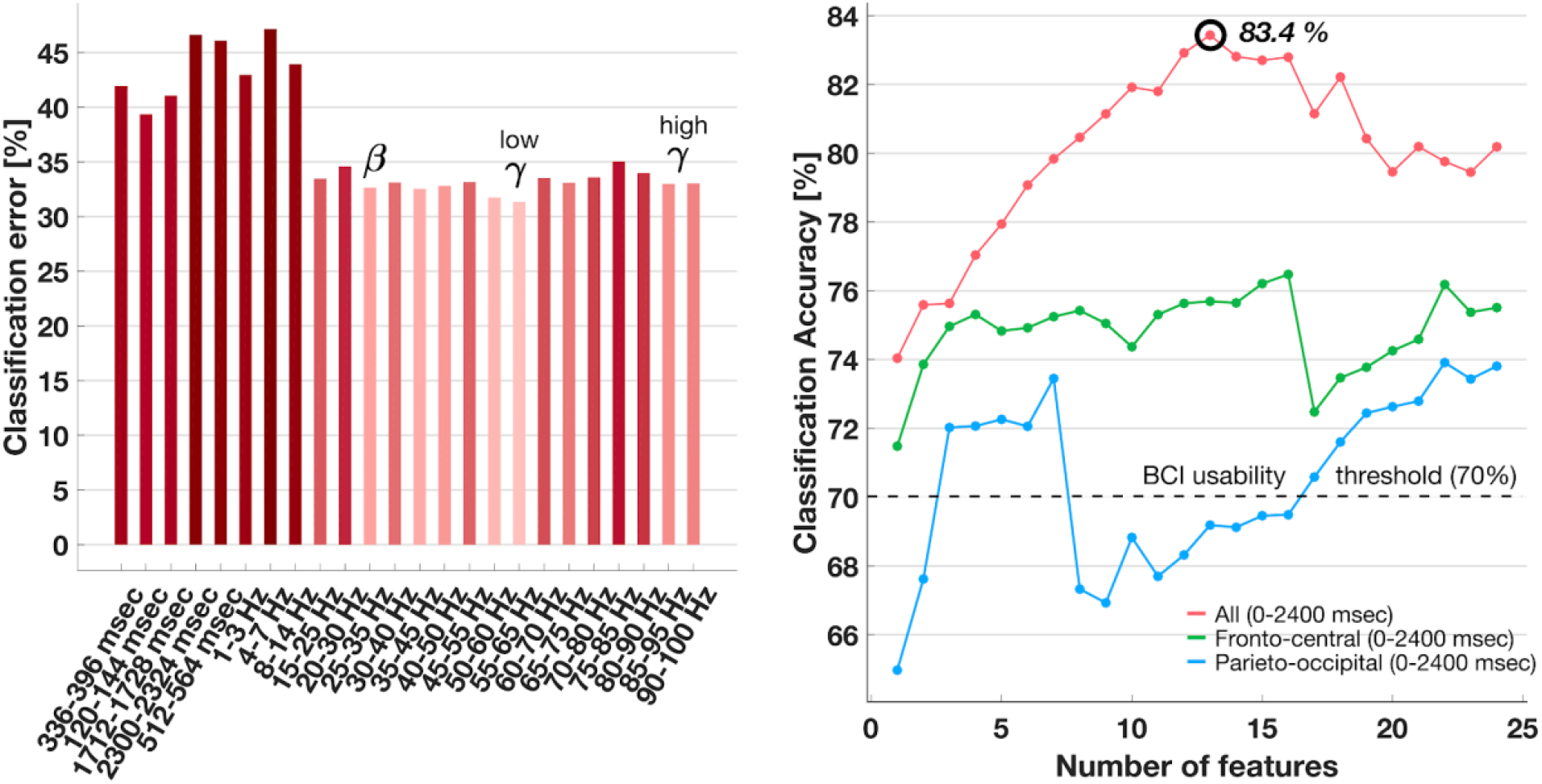
Spatio-temporal and Spectral classification of PC_OT/MW. LEFT: Averaged contribution to the classification error of each of the 5 spatiotemporal features and the 16 spectral features. Intensity of the colour denotes the contribution to the classification error with lighter colours corresponding to more discriminative features. RIGHT: Classification accuracies (averaged from 100 cross-validations) of iteratively added features ranked according to their discriminative power. The spectral features included in the classification are shown for the band power calculated over one interval (0 - 2400 msec) for three sets of channels (fronto-central, parieto-occipital and all channels). The x-axis depicts the number of features considered in the classification. The y-axis depicts the classification accuracy in percentage. The highest classification accuracy (83.4 %) was achieved with 13 features for all channels and spectral features extracted from the entire trial (0 - 2400 msec).

**Figure 4.**
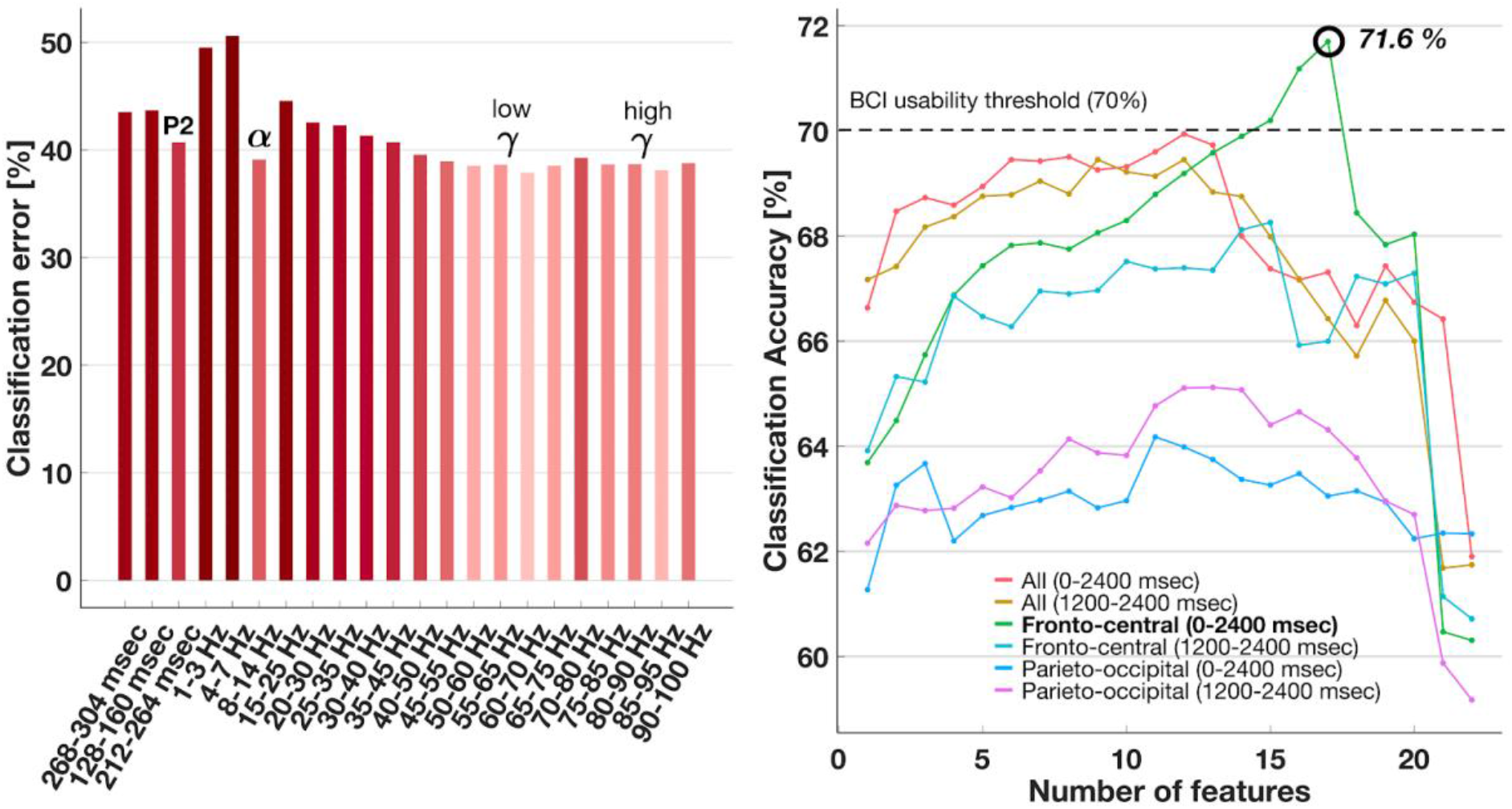
Spatio-temporal and Spectral classification of SC-dTUT/sTUT. LEFT: Averaged contribution to the classification error of each of the 3 spatiotemporal features and the 16 spectral features. Intensity of the colour denotes the contribution to the classification error with lighter colours corresponding to more discriminative features. RIGHT: Classification accuracies (averaged from 100 cross-validations) of iteratively added features ranked according to their discriminative power. The spectral features included in the classification are shown for the band power calculated over two intervals (0 - 2400 and 1200 2400 msec) for three sets of channels (fronto-central, parieto-occipital and all channels). The x-axis depicts the number of features considered in the classification. The y-axis depicts the classification accuracy in percentage. The highest classification accuracy (71.6%) was achieved with 17 features for the fronto-central channels and spectral features extracted from the entire trial (0 - 2400 msec).

**Figure 5.**
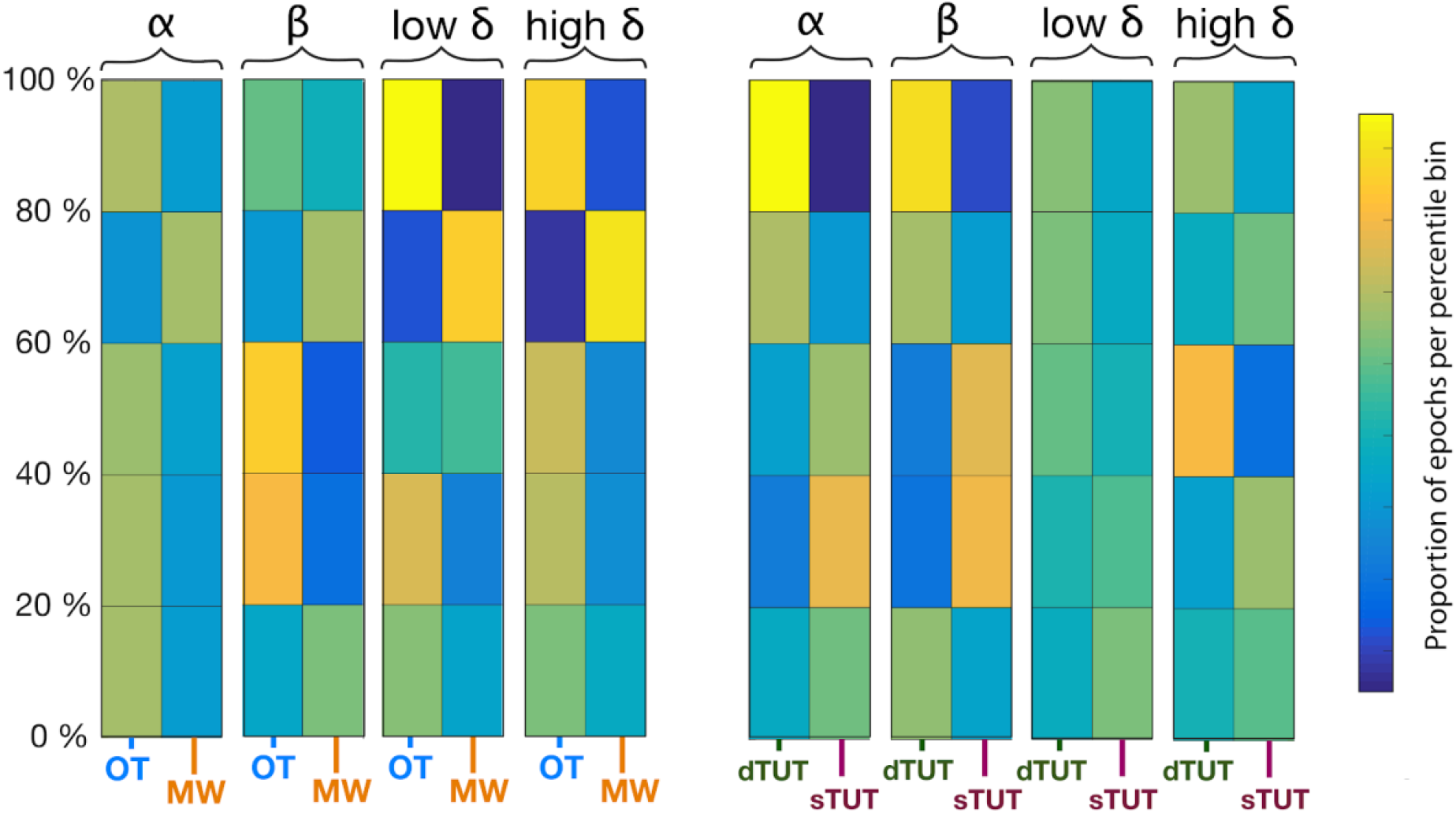
Bivariate histogram of the percentile distribution of epochs per band for all condition pairs. Each plot corresponds to a conditions-pair, PC_OT/MW (left) and SC_dTUT/sTUT (right). For each plot, the distribution of the number of epochs according to percentile bins (5 bins with each bin accounting for 20% of the distribution) is depicted per band. The colour of the tiles represent the number of epochs falling within a percentile bin with respect to band power, with higher percentile bins containing higher powered epochs. For example, for PC-OT/MW (left plot) the distribution for the low gamma band (50 -70 Hz) shows a proportionally high number of epochs with increased power belonging to the PC_OT condition. Conversely, the PC_MW condition had a proportionally smaller number of epochs with increased low gamma power. As a further example, SC_dTUT had a proportionally higher number of epochs with increased alpha activity relative to SC_sTUT.

In addition to gamma band power, beta power was found to be a discriminative feature for the PC_OT/MW classification (Figure 3).

Neural oscillations in the beta band has been found to be involved in several top-down processes such as functional inhibition and working memory encoding (Engel & Fries, 2010; Kilavik, Zaepffel, Brovelli, MacKay, & Riehle, 2013; Vázquez Marrufo, Vaquero, Cardoso, & Gómez, 2001). Beta has also been linked with task-related attention and attentional control ((Angelidis, van der Does, Schakel, & Putman, 2016; Laufs et al., 2006; Putman, Verkuil, Arias-Garcia, Pantazi, & van Schie, 2014; Ray & Cole, 1985) and reduced beta power was observed during off-task states (Braboszcz & Delorme, 2011; van Son et al., 2019). We found beta to be a discriminative feature for the PC_OT/MW classification (Figure 3) and a higher amount of high-powered beta epochs for dTUT, corroborating these prior studies. Our findings that both beta and gamma are characteristic of on-task and deliberate off-task states is in accord with the notion that the recruitment of top-down functions is necessary to uphold task-relevant or off-task patterns of cognition.

A central finding of our study is the discriminative power and prevalence of alpha activity with regard to periods when off-task thought is engaged in a deliberate manner. Oscillatory activity within the alpha range is thought to be related to attentional processes (Hanslmayr, Gross, Klimesch, & Shapiro, 2011) and in particular with those that relate to internal information (Cooper et al., 2003). One hypothesis is that alpha serves a regulator role between suppression and selection of information via a top down process inhibiting task-irrelevant cortical structures and corresponding sensory input (Foxe & Snyder, 2011; Klimesch, 2012; Mazaheri & Jensen, 2010). In the mind-wandering literature, alpha is a known electrophysiological correlate of off-task states (Baldwin et al., 2017; Boudewyn & Carter, 2018; Martel et al., 2014; O’Connell et al., 2009; Zhao, Wu, & Ou, 2013) and was recently found to be the most prominent oscillatory feature for the successful classification of off-task states (Jin et al., 2019). Given alpha’s putative role as supporting internally focused attention (Bazanova & Vernon, 2014; Benedek et al., 2014; Fink & Benedek, 2014) when engaged deliberately (Martel et al., 2019) our data provides further evidence that alpha is a general index of “internal attention” and suggests that it is a marker of deliberate off-task states in particular. Interestingly, besides the P2 component, spatio-temporal features exhibited the least discriminative ability, suggesting that spectral features could be sufficient for the implementation of a BCI able to identify off-task states. This is advantageous for the real-time classification based on ongoing EEG since endogenous brain signals, contrary to evoked activity, do not rely on external events, and thus, do not require the probing of an individual’s cognition with external stimuli (Berka et al., 2007; Blankertz et al., 2016). The results of our study highlight the importance of regarding off-task states as a heterogeneous class of mental states and confirm the feasibility of an EEG-based machine learning classifier able to detect different degrees of intentionality during off-task states. It should be noted that considering we found alpha activity to be idiosyncratic of intentional rather than spontaneous off-task states and that other studies ignoring this distinction observed it to be a marker of mind-wandering (Baldwin et al., 2017; Boudewyn & Carter, 2018; Jin et al., 2019; Macdonald et al., 2011; O’Connell et al., 2009; Zhao et al., 2013), it may be the case that at least some of the instances of mind-wandering measured in previous studies were of the deliberate kind.

However, a few cautionary remarks are in order when evaluating our findings. First, our classification was subject-independent, meaning that the classifier was trained on the entire datasets, without taking individual participants into account. Subject-independent classifiers might identify universal correlates to a specific brain state, thereby providing valuable information about the underlying neural mechanisms, and can be immediately applied on data from a new user. However, it would be good to note that taking into account subject-specific neural signatures would further improve the classification results of each subject.

And second, the univariate feature selection we used is suboptimal in the sense that despite being computationally efficient, only individual features were considered with respect to their usefulness and potential redundancies and reciprocities between features are ignored (Lotte, 2014). Multivariate algorithms for feature selection, on the other hand, evaluate features together, e.g. with mutual information (Peng, Long, & Ding, 2005). These methods have the advantage of considering redundancies and reciprocities between features, and obviate the need for manual selection based on a priori knowledge. Lastly, we did not conduct the search for discriminative features on a separate training set but rather on the entirety of the datasets for both condition-pairs. So, although the classification accuracy reported resulted from ten-fold cross-validations, the initial feature selection method might have contributed to an overestimation of the results and is a procedure that is not usable in an online scenario.

Notwithstanding these caveats, our approach was successful in its aims to determine whether the characteristic electrophysiological correlates of different aspects of off-task experiences could be reliably classified, and adds to the growing list of potential real-time markers of off-task thought. In the future, it might be of interest to resolve more dimensions of off-task states and establish idiosyncratic EEG profiles. A reliable EEG-based metric for the different aspects of mind-wandering would be instrumental in elucidating the temporal dynamics, neurocognitive processes and associated brain regions involved in the onset and maintenance of off-task states such as the switch between externally- and internally-directed attention. Future work could profitably examine whether, based on the present findings, a real-time prediction of off-task states can be achieved based on the spectral patterns of ongoing EEG dynamic.

